# The effect of pupil size and peripheral brightness on detection and discrimination performance

**DOI:** 10.1101/624783

**Authors:** Sebastiaan Mathôt, Yavor Ivanov

**Affiliations:** Department of Psychology, University of Groningen, The Netherlands; Research School of Behavioral and Cognitive Neurosciences, University of Groningen, The Netherlands

**Keywords:** pupillometry, pupil size, pupil light response, display polarity, display design, ergonomics, psychophysics

## Abstract

It is easier to read dark text on a bright background (positive polarity) than to read bright text on a dark background (negative polarity). This positive-polarity advantage is often linked to pupil size: A bright background induces small pupils, which in turn increases visual acuity. Here we report that pupil size, when manipulated through peripheral brightness, has qualitatively different effects on discrimination of fine stimuli in central vision and detection of faint stimuli in peripheral vision. Small pupils lead to improved discrimination performance, consistent with the positive-polarity advantage, but only for very small stimuli that are at the threshold of visual acuity. In contrast, large pupils lead to improved detection performance. These results are likely due to two pupil-size related factors: Small pupils increase visual acuity, which improves discrimination of fine stimuli; and large pupils increase light influx, which improves detection of faint stimuli. Light scatter is likely also a contributing factor: When a display is bright, light scatter creates a diffuse veil of retinal illumination that reduces image contrast, thus impairing detection performance. We further found that pupil size was larger during the detection task than during the discrimination task, even though both tasks were equally difficult and similar in visual input; this suggests that the pupil may automatically assume an optimal size for the current task. Our results may explain why pupils dilate in response to arousal: This may reflect an increased emphasis on detection of unpredictable danger, which is crucially important in many situations that are characterized by high levels of arousal. Finally, we discuss the implications of our results for the ergonomics of display design.

---

You are probably reading this text as dark letters on a bright background. And if not, then you might consider doing so, because it is easier to read dark letters on a bright background (positive polarity) than it is to read bright letters on a dark background (negative polarity). This positive-polarity advantage has been well-established in human-factors research (Buchner, Mayr, & Brandt, 2009; Dobres, Chahine, & Reimer, 2017; Piepenbrock, Mayr, & Buchner, 2014b, 2014a; Taptagaporn & Saito, 1990), and is often studied using proofreading experiments. For example, Piepenbrock et al. (2014b) asked participants to verbally report all misspelled words in a short text. The authors found that participants read faster, and spotted more mistakes, when the text was presented in a positive polarity, compared to a negative polarity. Findings such as these are among the reasons that most websites and word-processing software use a positive polarity.

The positive-polarity advantage is likely related to pupil size. When the background of a display is bright, the pupil constricts, compared to when the background is dark; this is the pupil light response (reviewed in Mathôt, 2018; Mathôt & Van der Stigchel, 2015). In terms of visual perception, there are three main consequences of a bright background and the resulting pupil constriction. The first consequence is negative: A bright background, as any source of brightness, results in light scatter; that is, some of the incoming light is not focused, but instead spreads over a large part of the retina. This results in a diffuse veil of light that reduces image contrast. The second consequence is also negative: Small pupils reduce the amount of light that falls on the retina, and thus reduce the signal-to-noise ratio of the image. The third consequence is positive: Small pupils suffer less from optical distortions that reduce image quality, and thus increase visual acuity (Campbell & Gregory, 1960; Liang & Williams, 1997; M. Lombardo & Lombardo, 2010; Woodhouse, 1975); that is, small pupils see sharper.

When reading text that is presented with sufficiently high contrast, as is typically the case in daily life, the advantage of increased visual acuity seems to outweigh the disadvantages of reduced signal-to-noise ratio and increased light scatter. Therefore, it is easier to read dark text on a bright background (when pupils are small), especially when the text is written in a small font (Piepenbrock et al., 2014a).

There is also some neurophysiological evidence that small pupils increase visual acuity. For example, Bombeke and colleagues (2016) manipulated pupil size by having participants covertly attend to either a bright or a dark disk, which respectively constricts or dilates the pupils, without changing eye position or visual input (cf. Binda, Pereverzeva, & Murray, 2013; Mathôt, van der Linden, Grainger, & Vitu, 2013). They then briefly presented a task-irrelevant but salient line-grating stimulus in peripheral vision, and measured the C1, an event-related potential (ERP) component that is associated with activity in primary visual cortex. The amplitude of the C1 was larger when participants attended the bright disk, compared to when they attended the dark disk. According to the authors, this result was due to the fact that small pupils improved the resolution of the C1-eliciting stimulus, in turn leading to a stronger neural response in primary visual cortex.

However, behavioral evidence for a link between pupil size and visual acuity is mixed. In a recent study by Ajasse, Benosman, and Lorenceau (2018), participants made a sequence of eye movements toward a configuration of disks; each disk had a different brightness, and the size of the pupil was therefore different depending on which disk the participant was fixating. While participants were fixating a disk, two gabor patches were briefly and sequentially presented in their visual periphery. The spatial frequency of the gabor patches differed, and participants indicated which of the two had the highest spatial frequency. The authors predicted that performance on this task should increase with decreasing pupil size (and thus with increasing brightness of the fixated disk). However, they found no such relationship; that is, performance did *not* depend on pupil size.

The results of Ajasse and colleagues (2018) show that small pupils do not lead to improved discrimination performance in every situation. Specifically, in their experiment, stimuli were presented in peripheral vision, where acuity is mostly limited by the reduced density of cone photoreceptors; therefore, in peripheral vision, optical blur due to large pupils likely has at most a very small effect on stimulus discrimination. However, the results of Bombeke and colleagues (2016), who also used a peripherally presented stimulus, suggest that under specific conditions a small-pupil advantage can also be found in peripheral vision.

In yet other situations, small pupils may even *impair* visual performance (reviewed in Mathôt, 2018; Mathôt & Van der Stigchel, 2015). Specifically, detecting faint stimuli in peripheral vision requires a high signal-to-noise ratio of the image, and visual acuity is only of secondary importance. In this case, large pupils may improve the signal-to-noise ratio of vision by increasing overall light influx. Therefore, stimulus detection in the visual periphery should benefit from large pupils. When large pupils are associated with a dark environment, as is typically the case in real life, this benefit should be even stronger, because the increased signal-to-noise ratio due to large pupils is accompanied by reduced light scatter due to the dark environment.

However, a study by Thigpen, Bradley, and Keil (2018) suggests that large pupils may not necessarily ‘boost’ neural responses to visual input. In their study, they presented a rapidly flickering stimulus, and measured so-called Steady-State Visually Evoked Potentials (ssVEPs): neural oscillations in visual cortex with the same frequency as the inducing stimulus. ssVEP power is believed to reflect the level of neural activity. Crucially, the authors found no relationship between ssVEP power and pupil size, and they interpreted this result as evidence for divisive normalization (Carandini & Heeger, 2012); that is, they suggested that visual responses, even in early visual cortex, are invariant to overall light influx and thus unaffected by pupil size.

Taken together, previous research has provided compelling evidence for an advantage of small pupils (and a bright background) for text reading (Buchner et al., 2009; Dobres et al., 2017; Piepenbrock et al., 2014b, 2014a; Taptagaporn & Saito, 1990). There is also some neurophysiological evidence for a small-pupil benefit for visual acuity (Bombeke et al., 2016; but see Ajasse et al., 2018). In contrast, there is no evidence for a large-pupil advantage for stimulus detection (e.g. Thigpen et al., 2018). Nevertheless, a large-pupil advantage for detection is clearly predicted based on the optical properties of the eye (see Mathôt, 2018; Mathôt & Van der Stigchel, 2015).

The aim of the current study is to demonstrate both a small-pupil advantage for discrimination of stimuli in central vision, and a large-pupil advantage for detection of stimuli in peripheral vision. We will manipulate pupil size by manipulating the brightness of the visual periphery, while presenting all task-relevant stimuli on a central gray disk of constant brightness.

## Experiments 1 and 2

The goal of Experiments 1 and 2 was to investigate whether pupil size, when manipulated through peripheral brightness, differentially affects performance on detection and discrimination tasks. In Experiment 1, participants detected, or discriminated the orientation of, a tilted Gabor patch. In Experiment 2, participants detected, or discriminated the lexicality of, a single word.

### Methods

#### Experiment 1

##### Participants, Ethics, and Apparatus

Nine naive observers participated in the experiment, after providing informed consent. The experiment was approved by the local ethics committee of Groningen University (16163-SP-NE and 16349-S-NE). Pupil size was recorded with an EyeLink 1000 (SR Research). Stimuli were presented with OpenSesame 3.1 (Mathôt, Schreij, & Theeuwes, 2012) on a 27” flat screen monitor with a resolution of 1920 × 1080 px.

##### Pupil-size manipulation

Pupil size was manipulated by varying the brightness of the visual periphery (low: 0.16 cd/m^2^, medium: 8.30 cd/m^2^, high: 52.26 cd/m^2^; see Figure 1), which corresponded to the full display (49.22° × 27.70°) except for a central gray disk. All task-relevant stimuli were presented on the central gray disk (2.84 cd/m^2^; diameter: 25.65°) that was kept constant throughout the experiment.

**Figure 1.**
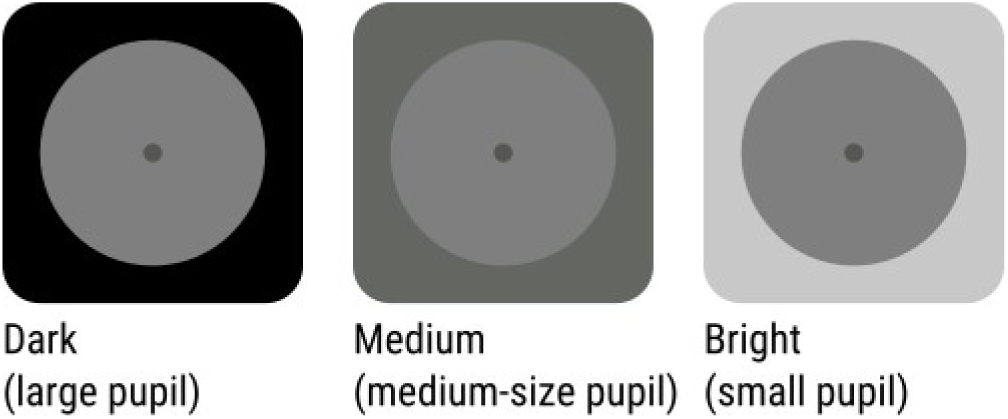
The luminance of the visual periphery was varied to manipulate pupil size. All task-relevant stimuli appeared on a central gray disk that was kept constant throughout the experiment.

##### Design

The experimental task (Discrimination or Detection) was varied between sessions. One experimental session consisted of five blocks.

The first two blocks of each session served to calibrate a Quest adaptive procedure, which varied the properties of the Target stimulus (see below) such that accuracy was kept at 75%. During these calibration blocks, peripheral brightness was set to 2.84 cd/m^2^. After these two blocks, the Quest procedure was stopped, and the Target was kept constant throughout the remainder of the session. Next, participants performed three blocks of 50 trials. Peripheral Brightness was varied between blocks (Figure 1).

Block order was fully counterbalanced, such that each possible order occurred once for each participant and task. Half the participants started on the first day with a Discrimination session followed by a Detection session, vice versa on the second day, etc. The other half of the participants started with a Detection session on the first day. In some cases, participants did more than two sessions per day. In total, participants performed 3,000 trials across 12 sessions in approximately six hours.

##### Discrimination Task

In the Discrimination task (see Figure 2a), each trial started with a central fixation dot (a uniform patch with a gaussian envelope with a standard deviation of 0.51° [20 px] and a peak brightness of 4.41 cd/m^2^) that was removed after 500 ms. After a random interval between 500 and 1,500 ms, drawn from a uniform distribution, a central Target Stimulus smoothly faded in and out over a period of 650 ms. The Target was a centrally presented sinusoidal grating with a gaussian envelope (a Gabor patch) with a standard deviation of 0.51° (20 px). To maintain accuracy at 75%, the spatial frequency, contrast, and orientation of the Target was varied with a Quest adaptive procedure during calibration blocks as described above.

**Figure 2.**
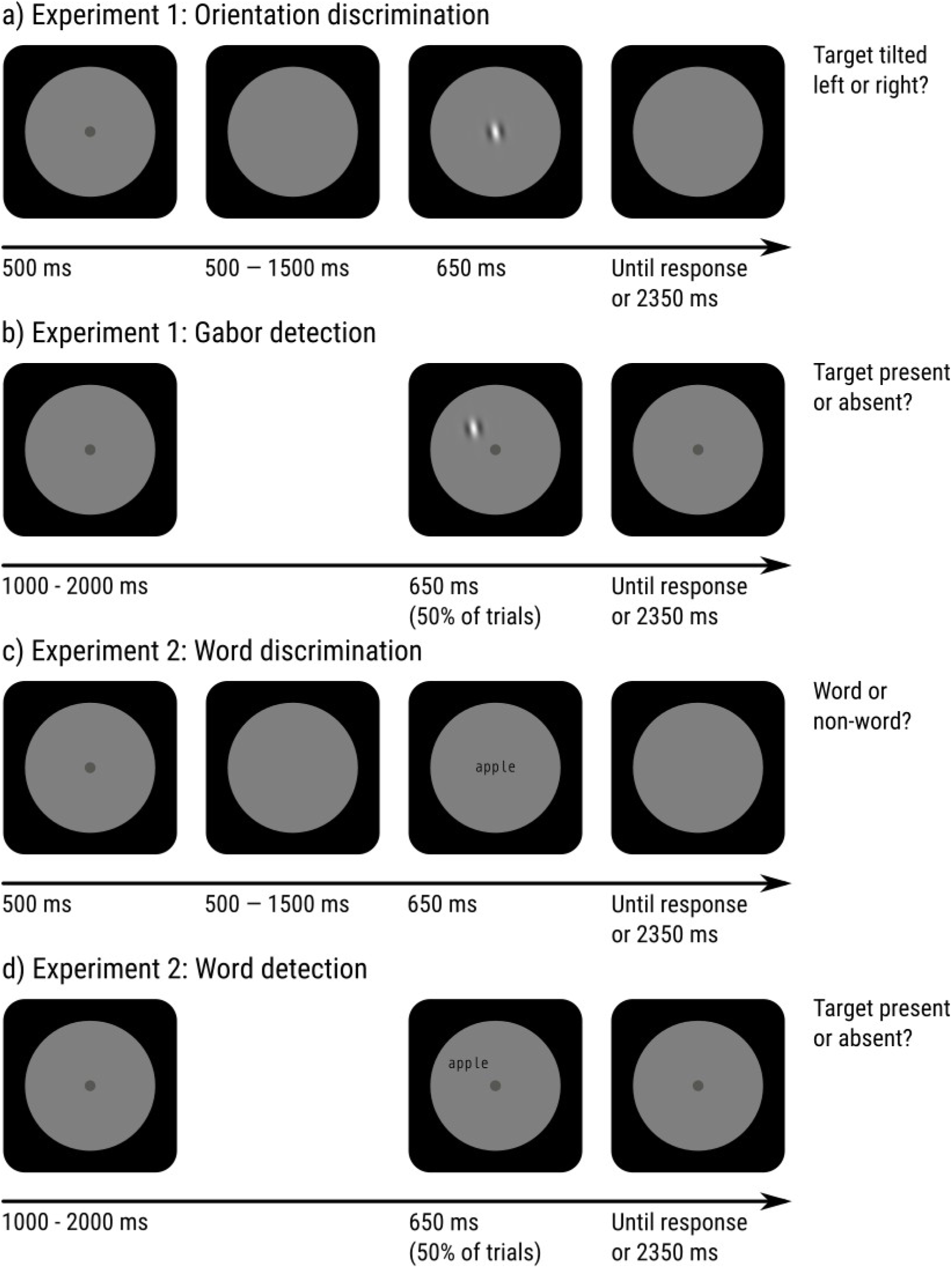
Schematic paradigm of Experiments 1 and 2. a) Orientation-discrimination task for Experiment 1. b) Orientation-detection task for Experiment 1. c) Word-discrimination (lexical decision) task for Experiment 2. d) Word-detection task for Experiment 2.

At any point during the trial, participants pressed the left arrow key if the Target was tilted counterclockwise from a vertical orientation, and the right arrow key if it was tilted clockwise. The trial ended 3 s after the onset of the Target.

##### Detection Task

In the Detection task (see Figure 2b), each trial started with a central fixation dot (4.41 cd/m^2^) that remained visible throughout the trial. On 50% of trials, after a random period drawn from a flat distribution between 1 and 2 s, a Target Stimulus was smoothly faded in and out over a period of 650 ms. The Target was identical to that of the Discrimination task, except that its standard deviation was 1.02° (40 px; i.e. twice as big), and that it was presented at a random point on an imaginary circle around the fixation dot with a radius of 7.70° (300 px).

At any point during the trial, participants pressed the space bar when they detected a Target, and did not press any key when they did not detect a Target. The trial ended 3 s after the onset of a Target (when present), or after a random interval between 4 and 5 s, drawn from a uniform distribution.

#### Experiment 2

Experiment 2 was in most ways identical to Experiment 1, and only the differences are described below.

##### Participants, Ethics, and Apparatus

Nine naive observers, most of whom had not participated in Experiment 1, participated in the experiment after providing informed consent. All participants were native Dutch speakers.

##### Stimulus selection

We selected the 750 most highly frequent words between four and six characters from the Dutch Lexicon Project (Keuleers, Diependaele, & Brysbaert, 2010), after manually (and based on our subjective impression) excluding overly offensive words. For each word, a matching pseudoword was generated with Wuggy (Keuleers & Brysbaert, 2010).

##### Design

Participants performed 1,500 trials across six sessions in approximately three hours. All participants saw all words and pseudowords once in a random order.

##### Task

Targets were (pseudo)words presented in a monospace font. In the Discrimination task, Targets were centrally presented, and participants pressed the left arrow key if the Target was a pseudoword and the right arrow key if it was a word (i.e. a lexical-decision task). In the Detection task, Targets were peripherally presented on 50% of trials, and participants pressed the space bar if they detected a target, and did not press any key otherwise. To maintain accuracy at 75%, the font size and contrast of the Target was varied.

### Results

We performed the same set of analyses on both experiments. The results from both experiments were very similar.

#### Task performance

To be able to directly compare performance in the Detection and Discrimination tasks, we used accuracy as our dependent measure. However, the results for the Detection task are similar when using *d’* (a measure of sensitivity that is based on signal-detection theory). Mean accuracy on the Detection task was 74.7% (Exp 1) and 73.2% (Exp 2). Mean accuracy on the Discrimination task was 74.5% (Exp 1) and 75.2% (Exp 2).

To test whether pupil size (as manipulated through peripheral brightness) affects performance (see Figure 3), and does so differently for the Discrimination and Detection tasks, we conducted a generalized linear mixed effects model (GLM) with Correct as dependent variable (binomial), Brightness (Low [reference], Medium, High), Condition (Detection [reference], Discrimination), and the Brightness × Condition interaction as fixed effects. We included only by-participant random intercepts, because more complex models failed to converge. (However, the results do not crucially depend on the exact model structure.) All mixed-effects analyses were conducted with the R package lme4 (Douglas et al., 2015).

**Figure 3.**
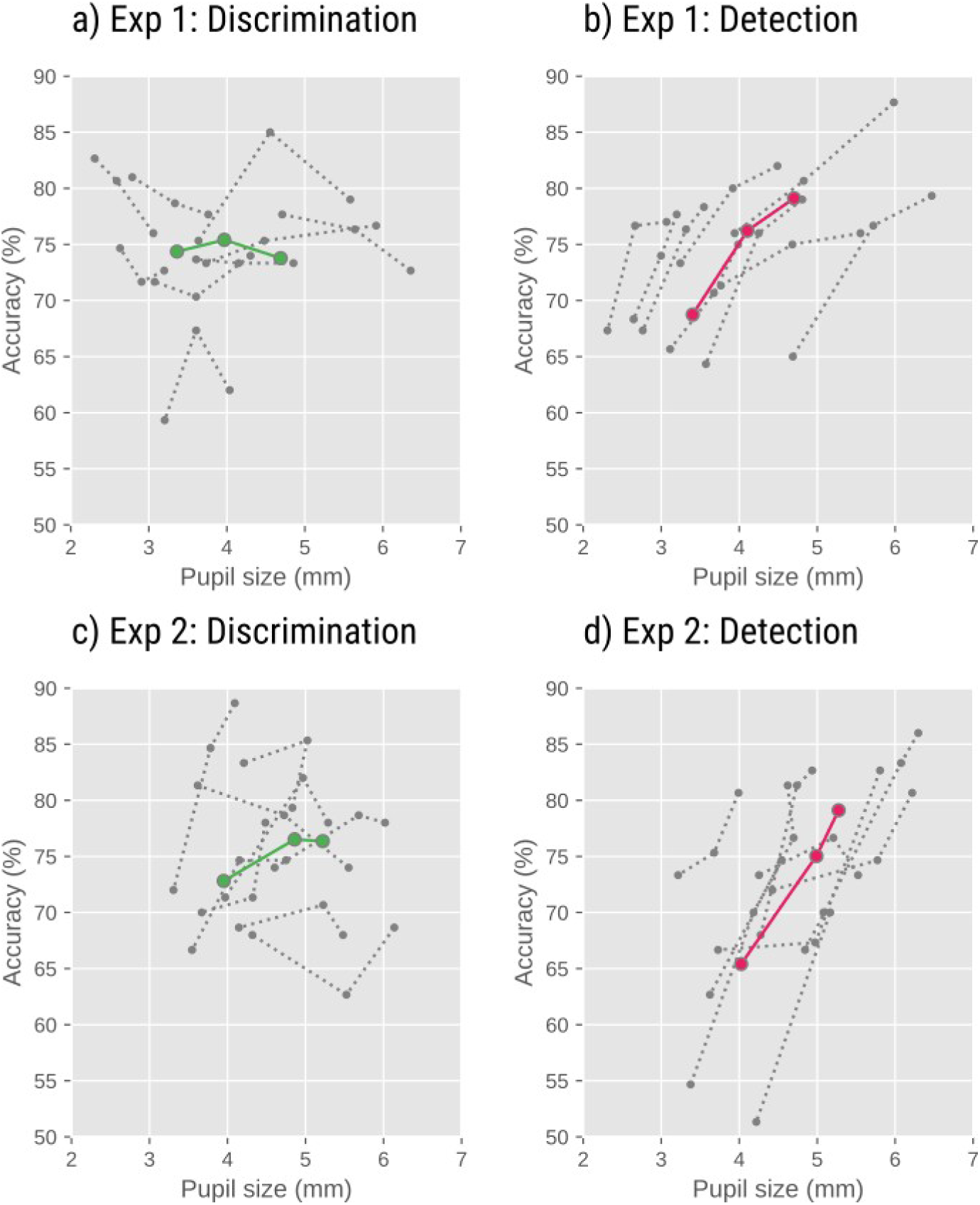
Detection accuracy increased with decreasing peripheral brightness, and thus increasing pupil size (b, d). However, there was no effect of peripheral brightness on discrimination performance (a, c). Gray dotted lines indicate individual participants. Colored solid lines indicate grand averages.

**Figure 4.**
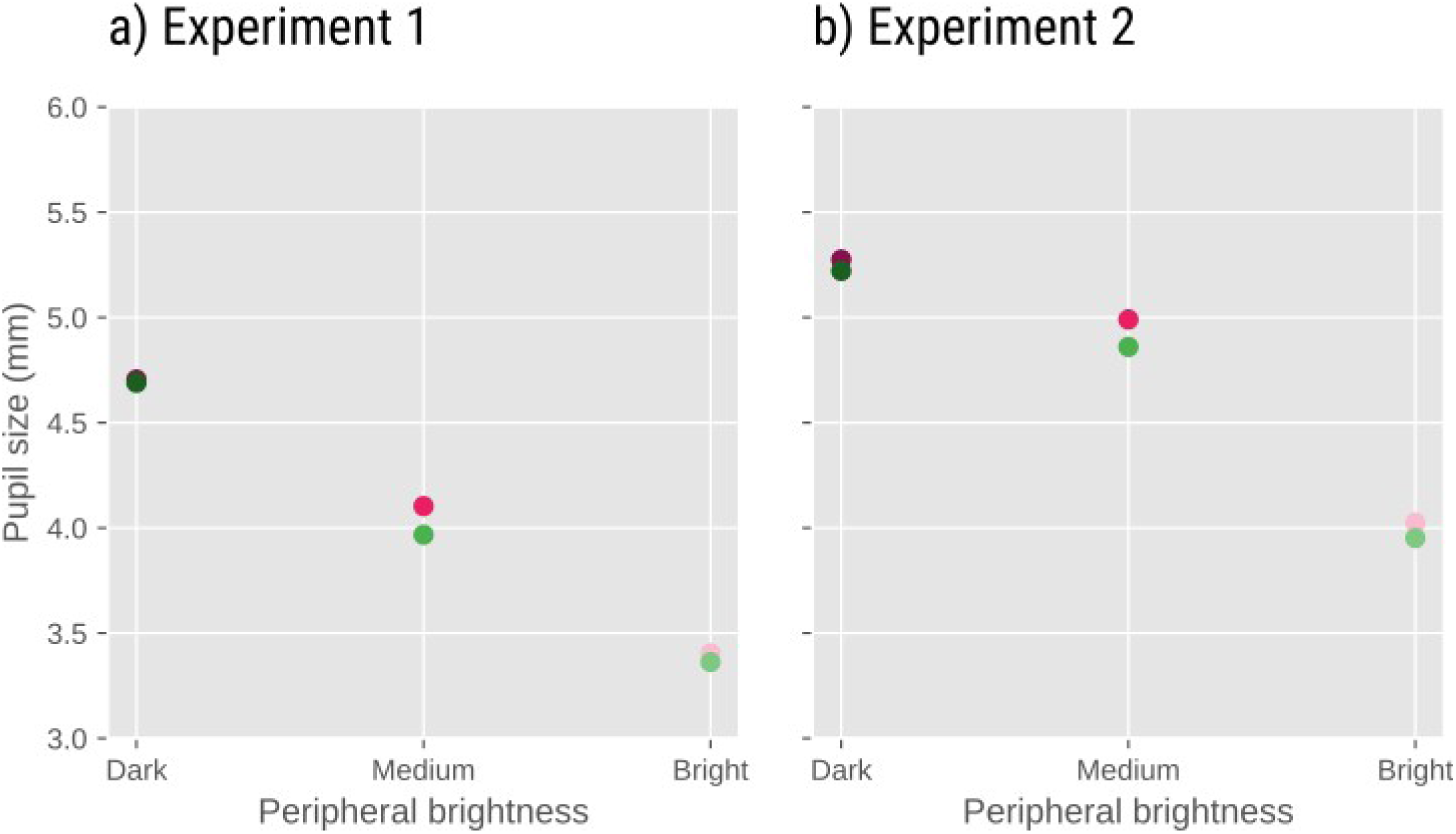
In both experiments, pupil size decreased with decreasing peripheral brightness. In addition, pupil size was slightly larger in the Detection than in the Discrimination condition.

There was an effect of Brightness (Exp 1: *Z* = −8.754, *p* < .001; Exp 2: *Z* = −8.005, *p* < .001), indicating that for the Detection (reference) Condition, accuracy decreased with increasing Brightness. There was an effect of Condition (Exp 1: *Z* = −5.343, *p* < .001; Exp 2: *Z* = −2.001, *p* = .045) indicating that for the Low (reference) Brightness, accuracy was lower for the Discrimination than Detection condition. Crucially, there was also a Brightness × Condition interaction (Exp 1: *Z* = 6.586, *p* < .001; Exp 2: *Z* = 4.114, *p* < .001), indicating that the effect of Brightness was driven by the Detection Condition, and not present in the Discrimination Condition.

To confirm this, we also analyzed the Discrimination Condition separately, in a model with only Brightness as fixed effect. Here we found no effect of Brightness in Exp 1 (*Z* = 0.503, *p* = .615), and only a weak effect of brightness in Exp 2 (*Z* = −2.153, *p* = 0.031).

#### Pupil size

The EyeLink provides pupil size in arbitrary units. To convert these units to millimeters of diameter, we first recorded artificial pupils (black circles printed on white paper) of different sizes, and then determined a function to convert EyeLink pupil units to pupil diameter (mm).

Mean pupil size during the Detection task was 4.1 mm (Exp 1) and 4.8 mm (Exp 2). Mean pupil size on the Discrimination task was 4.0 mm (Exp 1) and 4.7 mm (Exp 2).

Our brightness manipulation should have a large effect on pupil size. It is also possible that the task affects pupil size, despite the fact that the two tasks were equally difficult. To test this, we conducted a linear mixed-effects analysis (LMER) with Pupil Size as dependent measure and Brightness, Condition, and a Brightness × Condition interaction as fixed effects. Again, we included only by-participant random intercepts, because more complex models failed to converge.

There was an effect of Brightness (Exp 1: *t* = −87.405; *p* < .001 Exp 2: *t* = −58.165, *p* < .001), reflecting that pupil size decreased with increasing brightness. There was also an effect of Condition (Exp 1: *t* = −3.849, *p* < .001; Exp. 2: *t* = −4.340, *p* < .001), reflecting that pupil size was larger in the Detection than in the Discrimination condition. There was no notable Brightness × Condition interaction (Exp 1: *t* = −0.930, *p* = 0.352; Exp 2: *t* = −0.216, *p* = 0.829).

### Discussion

In summary, we found that detection performance was better with large pupils (and a dark periphery) than with small pupils (and a bright periphery). This effect was large, robust, and in the direction that we predicted. However, and unlike we predicted, we did not find that discrimination performance increased with decreasing pupil size (and thus increasing peripheral brightness); in fact, there was a slight effect in the opposite direction for Exp. 2.

In addition, we found that pupil size was larger in the Detection than in the Discrimination condition, even though both tasks were equally difficult.

A limitation of our setup for measuring discrimination performance was that we could not present very small stimuli: When presented at full contrast, even the finest possible grating (i.e. 2 px/cycle) or the smallest possible letter (5 × 5 pixels) could be discriminated without too much trouble by someone with normal vision. Therefore, to increase the difficulty of the discrimination task, we also reduced the contrast of the target stimulus, and our discrimination task was therefore not a pure measure of discrimination performance (a limitation that we addressed in Experiment 3).

## Experiment 3

In Experiment 3, we used a setup that allowed us to present very small letters. The aim of this experiment was to investigate whether we could observe an advantage of small pupils (and increased peripheral brightness) on discrimination performance in a task that tested the limits of visual acuity. If so, this would suggest that the absence of a small-pupil benefit in Experiments 1 and 2 was due to the fact that, in these experiments, our stimuli were not sufficiently fine to test the limits of visual acuity.

### Methods

#### Participants, Ethics, and Apparatus

20 naive observers participated in the experiment, after providing informed consent. The experiment was approved by the local ethics committee of Groningen University (16163-SP-NE and 16349-S-NE). Pupil size was recorded with an EyeLink 1000 (SR Research). Stimuli were presented with OpenSesame 3.1 (Mathôt et al., 2012) on three separate 7” tablets (Samsung Galaxy Tab 7), each with a resolution of 1280 × 800 px. Two tablets were presented nearby, on both sides, of the participant’s head, and served as light sources. One tablet was placed in front of the participant, at a distance of 5.5 m, and served as the target display.

#### Task and Design

Each trial started with the presentation of a black central fixation dot (R=0.11° [90 px]) for 500 ms on the target display. Next, a lowercase or uppercase letter (‘a’, ‘b’, ‘d’, ‘e’, ‘g’, ‘h’, ‘l’, ‘m’, ‘n’, ‘q’, ‘r’, or ‘t’) was presented for 2.5 s. Letters were presented centrally in black monospace font. The size of the letters was varied with a Quest adaptive procedure to converge on 75% accuracy. Participants pressed the ‘z’ key to indicate that the letter was lowercase, and the ‘/’ key to indicate that the letter was uppercase.

The experiment started with a practice block of 75 trials during which the background of all tablets was gray (72.15 cd/m^2^ for target display; 69.01, 64.70 cd/m^2^ for peripheral displays). This practice block also served to determine an appropriate font size to start with during the experimental blocks. Next, participants performed six experimental blocks during which the brightness of the light-source tablets was varied (343.30 cd/m^2^ and 345.20 cd/m^2^ for the two peripheral displays], medium [69.01 cd/m^2^, 64.70 cd/m^2^], or dark [0.83 cd/m^2^, 0.88 cd/m^2^]) while the background of the target display remained gray. Each experimental block started with the font size that the practice block had ended with. The order of the experimental blocks followed a counterbalanced ABCABC design. In total, participants performed 375 trials in approximately 40 minutes.

### Results

#### Task performance

For each participant and Brightness Level separately, we took the final font size of each block as a measure of performance (because the Quest procedure varied font size depending on the participant’s performance).

To test whether peripheral brightness (and pupil size) affects performance, we conducted a linear mixed effects model (LME) with Final Quest Value (on which font size was based) as dependent measure and Brightness as fixed effect. We included by-participant random intercepts and slopes. There was an effect of Brightness (*t* = −2.356, *p* = .029), indicating that discrimination performance increased with increasing peripheral brightness (Figure 6).

**Figure 5.**
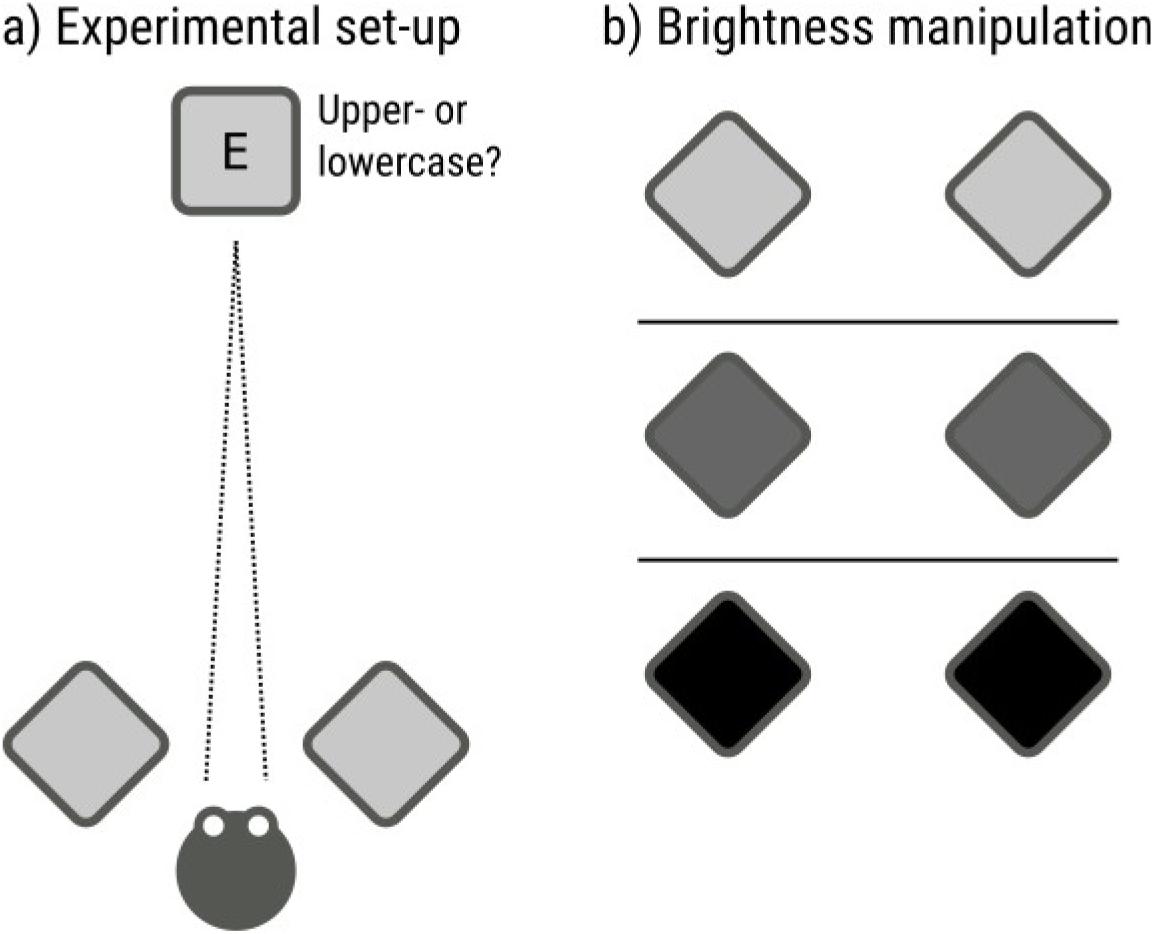
Schematic set-up and paradigm of Experiment 3. a) Participants indicated whether a target letter was uppercase or lowercase. b) Pupil size was manipulated by varying the brightness of two displays that were positioned near the participant, and flanked the target display.

**Figure 6.**
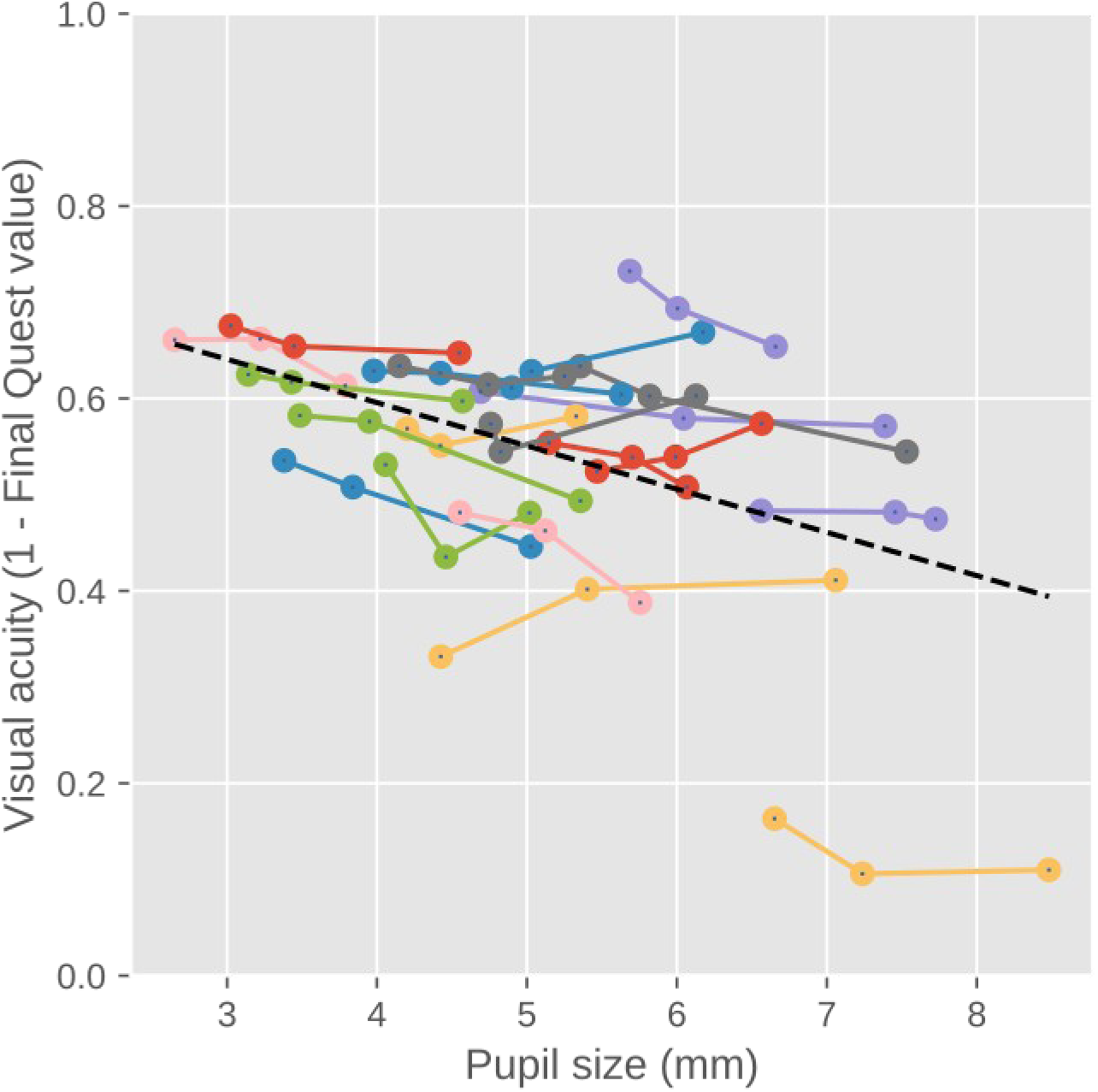
Discrimination performance increased with increasing peripheral brightness (and decreasing pupil size). Lines correspond to individual participants. Dots correspond to different levels of peripheral brightness, such that the highest peripheral brightness corresponds to the smallest pupil size. In addition, participants with smaller overall pupils had higher discrimination performance.

To test whether individual differences in pupil size also affect performance, we determined, for each participant separately, the mean pupil size during the entire experiment, and the mean Quest value during the last half of all blocks. There was a clear correlation between the two measures (*r* = -.508, *p* = .022), indicating that participants with smaller overall pupils had higher performance.

#### Pupil size

To test whether our brightness manipulation affects pupil size, we conducted an LME with Pupil Size as dependent measure and Brightness as fixed effect. We included by-participant random intercepts and slopes. There was an effect of Brightness (*t* = −12.662, *p* < .001), indicating that pupil size decreased with increasing brightness of the flanking tablets. Pupil size was converted from arbitrary units to millimeters of diameter with the same procedure as used for Experiments 1 and 2.

### Discussion

As predicted, we found that small pupils (and increased peripheral brightness) improved discrimination performance. In, addition we found that participants with small pupils had higher discrimination performance. That is, there was a clear link between pupil size and discrimination performance, both when pupil size was manipulated experimentally, and when considering individual differences.

## General Discussion

Here we report that pupil size, when manipulated through peripheral brightness, has qualitatively different effects on discrimination of fine stimuli in central vision and detection of faint stimuli in peripheral vision.

Specifically, we found that small pupils (and thus a bright periphery) lead to improved discrimination of small letters presented in central vision. This is consistent with previous studies that showed a so-called positive-polarity advantage; that is, it is easier to read dark letters on a bright background (positive polarity) than it is to read bright letters on dark background (negative polarity) (Buchner et al., 2009; Dobres et al., 2017; Piepenbrock et al., 2014b, 2014a; Taptagaporn & Saito, 1990). We observed this effect only (but highly reliably) with very small letters that were at the limits of visual acuity. This is consistent with a previous study showing that the positive-polarity advantage is most pronounced for small letters (Piepenbrock et al., 2014a). The small-pupil benefit for discrimination is likely due to the fact that visual acuity is highest with small pupils, which suffer less from optical distortions that reduce visual acuity (Campbell & Gregory, 1960; Liang & Williams, 1997; M. Lombardo & Lombardo, 2010; Woodhouse, 1975; Bombeke et al., 2016).

We further found that large pupils (and thus a dark periphery) improved detection of faint stimuli that were presented at an unpredictable location in peripheral vision. This large-pupil benefit for detection is likely due to two factors. First, large pupils increase light influx, which increases the signal-to-noise ratio of vision, which in turn facilitates detection of very faint stimuli (but perhaps not, or hardly, of stimuli that are presented well above the detection threshold, as used for example by Thigpen et al., [2018]). Second, the dark periphery that we used to induce large pupils resulted in reduced light scatter (M. Lombardo & Lombardo, 2010), in turn resulting in increased image contrast, thus making it easier to detect stimuli. Therefore, reduced light scatter likely also contributed to the large-pupil benefit (which is therefore in part likely a dark-periphery benefit).

Our results offer a possible explanation for why the pupils dilate in response to increased arousal (e.g. Mathôt, 2018; Mathôt & Van der Stigchel, 2015). Situations that require fine discrimination are often characterized by low levels of arousal, and situations that require detection are often characterized by high levels of arousal. For example, arousal is low when a person is reading a book, or when an animal is foraging for food. In such cases, it is crucially important to identify what you’re looking at. In contrast, arousal is high when a person is afraid, or when an animal is on the lookout for predators. In such cases, it is crucially important to detect unexpected dangers. In other words, pupil dilation in response to arousal may reflect an increased emphasis on visual sensitivity, at the expense of visual acuity, to meet the demands of the situation.

An incidental yet striking result is that pupil size was larger during the detection task than during the discrimination task. Because there was no systematic difference in difficulty between the two tasks, this pupil-size difference is likely not due to differences in mental effort (which is known to affect pupil size, see e.g. Mathôt, 2018). One possibility is that, in the detection task, the pupil dilated as a result of attention being directed to peripheral rather than central vision (cf. Brocher, Harbecke, Graf, Memmert, & Hüttermann, 2018; Daniels, Nichols, Seifert, & Hock, 2012). An even more interesting possibility is that the pupil automatically assumes a size that is optimal for the current task, and that arousal-related pupil responses are merely one example of this general principle.

Our results also have implications for the ergonomics of display design. First, as was already well-established, visual information that requires fine discrimination, such as text, is best displayed on a bright background (positive polarity), which induces small pupils (Buchner et al., 2009; Dobres et al., 2017; Piepenbrock et al., 2014b, 2014a; Taptagaporn & Saito, 1990). Second, and this is a novel insight that results from our findings, visual information that should capture attention, such as notifications, may be best displayed on a dark background (negative polarity), which induces large pupils. Importantly, pupil size is determined by the overall level of display brightness, although with a bias towards central vision (e.g. Crawford, 1936). Therefore, there is no point in presenting notifications in a dark region of a display that is otherwise bright. This poses a dilemma when designing displays that contain text as well as notifications. In such cases, the relative importance of the different kinds of information should determine whether the display should use a positive or a negative polarity.

In summary, we have shown that small pupils, induced through a bright periphery, lead to improved discrimination of fine stimuli in central vision. In contrast, large pupils, induced through a dark periphery, lead to improved detection of faint stimuli in peripheral vision.

## Materials and availability

Data and experimental materials can found at https://osf.io/h389s/

## Notes

https://osf.io/h389s/

